# Overt visual attention modulates decision-related signals in the frontal cortex

**DOI:** 10.1101/2024.10.25.620227

**Authors:** Blair R. K. Shevlin, Rachael Gwinn, Aidan Makwana, Ian Krajbich

## Abstract

When indicating a preference between two options, decision makers are thought to compare and accumulate evidence in an attention-guided process. Little is known about this process’s neural substrates or how visual attention affects the representations of accumulated evidence. We conducted a simultaneous eye-tracking and fMRI experiment in which human subjects gradually learned about the value of two food-lotteries. With this design we were able to extend decisions over a prolonged time-course, manipulate the temporal onset of evidence, and therefore dissociate sampled and accumulated evidence. We observed inconsistent correlations of both sampled and accumulated evidence with activity in the ventromedial prefrontal cortex (vmPFC), the ventral striatum, and the intraparietal sulcus (IPS), and more consistent correlations of accumulated evidence with activity in the dorsolateral prefrontal cortex (dlPFC) and pre-supplementary motor area (pre-SMA). We also found that more gaze on an option increased its choice probability and that gaze consistently amplified accumulated-value signals above and beyond the non-gaze-modulated signals in the pre-SMA and partially in the dlPFC, providing novel evidence that visual attention has lasting effects on decision variables and suggesting that activity in the pre-SMA and dlPFC reflect gaze-weighted accumulated evidence. These results shed new light on the neural mechanisms underlying gaze-driven decision processes.

## Introduction

Many decisions involve a process of value computation and comparison between options. Imagine you are planning a vacation based on online travel-guides describing two locations. As you research these options, you sample pieces of information that support one option or another to varying degrees. For example, one destination might have cheaper flights while the other has cheaper hotels. Additionally, some information may attract more of your attention, such as pictures of waterfalls or ancient ruins. As you evaluate your options, you must integrate these pieces of information and eventually decide when to stop and make a final decision.

Decisions like this are thought to rely on a bounded, evidence-accumulation process that depends on factors such as the value of the sampled information and shifts in attention. According to this framework, when two options are similar in value, evidence accumulates more slowly towards the decision threshold, resulting in longer response times (RT) and more opportunity for shifts in attention to influence the choice outcome. In contrast, when one option is clearly superior, evidence accumulates more rapidly and the decision is made quickly with less of a relation between gaze and choice. This choice process produces reliable, quantitative patterns in choice, RT, and eye-tracking data (Ashby et al., 2016; Callaway et al., 2021; Gluth et al., 2018; Krajbich et al., 2010; Smith & Krajbich, 2018). For instance, decisions with similar values are more random (i.e., less predictable), tend to take more time (Konovalov & Krajbich, 2019), and can be experimentally manipulated by diverting attention towards one option more than the other (Bhatnagar & Orquin, 2022; Gwinn et al., 2019; Pärnamets et al., 2015; Pleskac et al., 2022; Tavares et al., 2017). Critically, these behavioral measures do not simply correlate; rather, they exhibit precise quantitative relationships consistent with evidence accumulation models (Konovalov & Krajbich, 2019).

Sequential sampling models (SSMs) offer a framework to understand this evidence accumulation-driven choice process. SSMs vary in specific details, but generally share some core features. When faced with a decision, people begin to sample information in favor of each option. In many models, this information is evaluated and converted into relative evidence for one option or the other. Relative evidence builds up over time until there is enough to commit to an option (Busemeyer & Townsend, 1993; Ratcliff & Smith, 2004).

Most SSMs can be broken down into two key components: inputs and integrators (Bogacz et al., 2006). Inputs encode the drift rate – the rate of evidence accumulation – and integrators encode the decision variable – the amount of accumulated evidence.

Each option has its own input. In the context of value-based decisions, the input represents the value of the currently considered piece of information. For a given option, the average input value is generally assumed to be constant over the course of the decision but does vary randomly from one instant to the next due to stochasticity in the sampling process (Shadlen & Shohamy, 2016).

Integrators accumulate the stochastic sequences of sampled input values. Typically, each option has its own integrator (Busemeyer & Townsend, 1993; Gold & Shadlen, 2007; Krajbich & Rangel, 2011; Usher & McClelland, 2001; Wang, 2002) which accumulates the evidence from that option’s input, but is also inhibited by the other options’ inputs or integrators. Thus, each integrator represents the accumulated, relative evidence favoring a given option. Once one of the integrators’ accumulated evidence reaches a pre-determined threshold, the corresponding option is chosen. In contrast to the input values, these accumulated values dynamically evolve over the course of the decision.

The neural inputs and accumulated values have been successfully identified in perceptual decision making (for recent reviews see: Forstmann et al., 2016; Hanks & Summerfield, 2017; O’Connell et al., 2018; Ratcliff et al., 2016), but it has proven more challenging for value-based decisions. The main reason is that decisions are typically very quick, with RTs shorter than the time resolution of functional magnetic resonance imaging (fMRI) (but see Gluth et al., 2012).

Evidence for the neural substrates of value-based SSMs have typically come as trial-level measures correlating with model parameters (Hare et al., 2011; Rodriguez et al., 2015), scalp-level electric activity from electroencephalography (EEG) (Polanía et al., 2014), or a combination of the two (Pisauro et al., 2017). Taken together, these studies have implicated a fronto-parietal network underlying value-based decision-making, with more ventral/frontal regions serving as inputs (for reviews see: Bartra et al., 2013; Clithero, 2018) and more dorsal/parietal regions serving as integrators.

Despite their many strengths, these past value-based experiments have been limited by their inability to determine whether purported integrator regions are accumulating evidence or instead representing unchosen values (Boorman et al., 2011; Kolling et al., 2016; Wittmann et al., 2016), decision conflict (Frömer et al., 2019; Kaanders et al., 2021; Shenhav et al., 2014, 2016; Vassena et al., 2020), or time on task (Grinband et al., 2011; Holroyd et al., 2018; Mumford et al., 2024). In most experiments, these variables are highly correlated and difficult to distinguish. Increasing the value of the worse option while holding the better option constant will simultaneously increase the perceived conflict, increase the deliberation time, and slow the rate of evidence accumulation.

To distinguish between accumulated evidence and the other confounding explanations, we sought a factor that modulates accumulated evidence within a decision, independent of time. For this we turned to visual attention, measured with eye-tracking. Research on the attentional drift diffusion model (aDDM) has argued that gaze amplifies value during the choice process (Bhatnagar & Orquin, 2022; Krajbich et al., 2010; Sepulveda et al., 2020; Smith & Krajbich, 2019; Westbrook et al., 2020). This means that current gaze location should amplify value signals in the input regions, and that the balance of gaze allocation over the course of the decision should amplify accumulated evidence signals in the integrator regions. Both human and non-human primate research has confirmed gaze effects on value inputs in the orbitofrontal cortex (Lim et al., 2011; McGinty et al., 2016; Rich & Wallis, 2016; Hunt et al., 2018; but see McGinty, 2019). However, it has yet to be shown that these gaze-modulated inputs are integrated into accumulated decision values. Gaze modulated signals in purported integration regions would provide critical evidence against the alternative explanations (i.e., conflict, time, or unchosen value).

Here, we present the results of an fMRI experiment designed to provide evidence that integrator regions accumulate gaze-weighted evidence. Our approach was to slow down the decision process by gradually presenting choice-relevant information. Our task design allowed us to extend the decision-making period to approximately a minute, while also allowing us to dissociate the inputs’ sampled value (SV) signals from the integrators’ accumulated value (AV) signals (Gwinn et al., 2019). The inputs represent the perceived value of stimuli currently on the screen, while the integrators represent the values of previously presented stimuli within that choice problem. We simultaneously collected eye-tracking data, allowing us to test whether gaze modulates SV and AV representations. We specifically focus on the difference in sampled values between the left and right options (Δ*SV*) and the corresponding difference in accumulated values between the left and right options (Δ*AV*). We hypothesized that we would find (1) a positive correlation between gaze-weighted |Δ*SV*| and activity in the reward network (the ventromedial prefrontal cortex (vmPFC) and ventral striatum), and (2) a positive correlation between gaze-weighted |Δ*AV*| in the pre-supplementary motor area (pre-SMA) (Aquino et al., 2023), dorsolateral prefrontal cortex (dlPFC), and intraparietal sulcus (IPS).

## Results

### Experiment description

Our choice task builds on an extensive literature examining choices between familiar snack foods. Instead of choosing between two food items, which typically only takes a few seconds, we asked subjects to choose between two food lotteries. A lottery consisted of 3-6 different items, each with a different probability of being selected. Subjects did not know anything else about the lotteries; they had to learn about them from experience (Hertwig et al., 2004). Specifically, subjects sampled a random draw from both lotteries every 4-8 seconds. They continued to sample random draws until they were ready to stop and choose one of the two lotteries (Fig. 1). Choosing a lottery led to a final random draw from that lottery, revealing the actual food that the subject would receive if that trial was rewarded at the end of the experiment.

**Figure 1.**
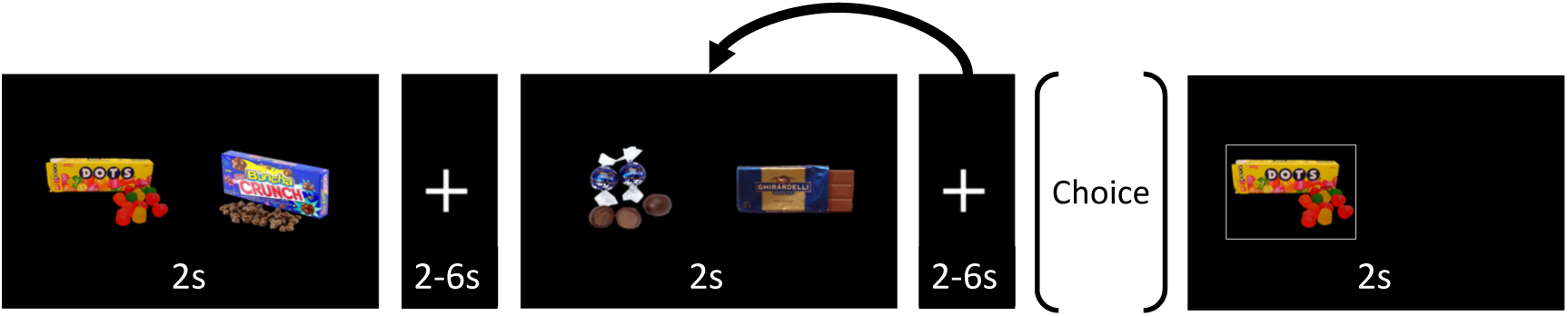
Task timeline. Subjects chose between two snack food lotteries on each trial. Subjects learned about the lotteries through random food draws. Every 4-8 seconds, subjects sampled a new draw from each lottery. They were allowed to sample as many times as they wanted but were incentivized to sample approximately 7 draws per trial. Sampled food draws were presented for 2 seconds, followed by a fixation cross appearing for 2-6 seconds with random jitter. The trial ended when the subject chose the left or right lottery, using the respective index finger. Upon making their choice, subjects were presented with a food drawn from their chosen lottery.

We placed no explicit limit on the number of draws subjects could sample within each trial (i.e. pair of lotteries). To prevent subjects from spending the entire session on a single trial, we gave them 45 minutes to make at least 60 choices. Subjects were informed that any unmade choices would be randomly completed by the computer, and any trials beyond 60 would be added to the list from which the rewarded trial would be drawn. If a subject were to sample the same number of stimuli in each trial, the optimal number would be seven. However, subjects could (and did) vary their number of samples throughout the experiment; we observed substantial variability in the number of samples per trial within most of our subjects (mean number of samples = 6.37 and mean within-subject *SD* = 2.61).

### Choice

We constructed each trial’s sequence of items pseudo-randomly to minimize the correlation between |Δ*SV|* (i.e., the absolute difference between the left and right values of the currently presented stimuli) and |Δ*AV*| (i.e., the absolute sum of Δ*SV* at a given time within a trial) (see Methods). For the first sample in each trial, Δ*SV* and Δ*AV* were always equal. After subsequent samples, Δ*SV* diverged from Δ*AV*, yielding two distinct time courses to look for in the fMRI data (Fig. 2).

**Figure 2.**
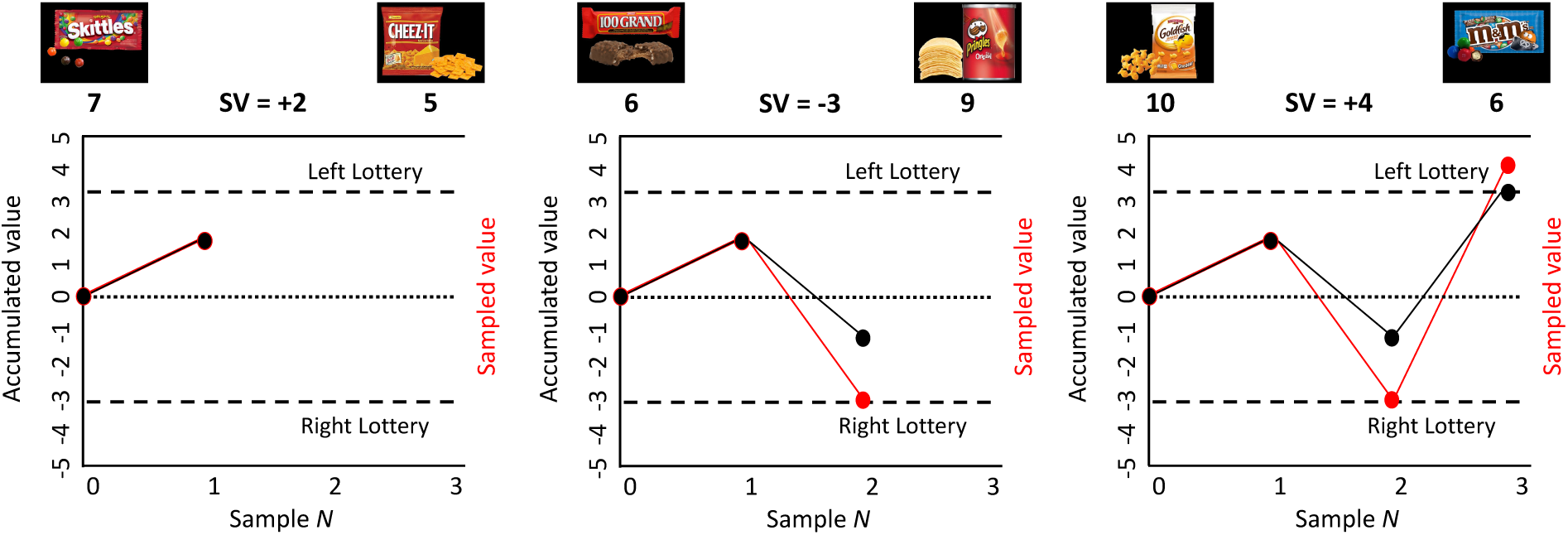
Example trial with the sampled value and accumulated value. The sampled value (Δ*SV*; red) and accumulated value (Δ*AV*; black) are plotted for this example trial. For the first draw, Δ*SV* and Δ*AV* are identical. However, as the trial proceeds, the two signals diverge. In the model, a choice is made when the |Δ*AV|* reaches a pre-specified decision boundary.

To measure the subjective value of each stimulus, we separately asked subjects to rate all the food items. Before entering the scanner, subjects rated 148 unique food items based on how much they would like to eat them at the end of the experiment. These ratings were incentivized (see Methods) and we retained only the positively rated items (0 to 10) for the choice task. We used each subject’s ratings to calculate SV and AV.

### Behavioral results

A core assumption of SSMs is that individuals decide based on the evidence accumulated over the course of the decision. We thus anticipated that subjects would choose in line with Δ*AV* and not just the most recent Δ*SV* in the trial. We tested this key assumption with a mixed-effects logistic regression of choosing the left lottery on Δ*SV* and Δ*AV* at the time of choice. Subjects chose in line with both (Δ*AV* excluding the final samples: *β* = 0.062, *SE*= 0.010, *p* < 0.001; Δ*SV*: *β* = 0.257, *SE*= 0.024, *p* < 0.001) (Fig. 3A).

**Figure 3.**
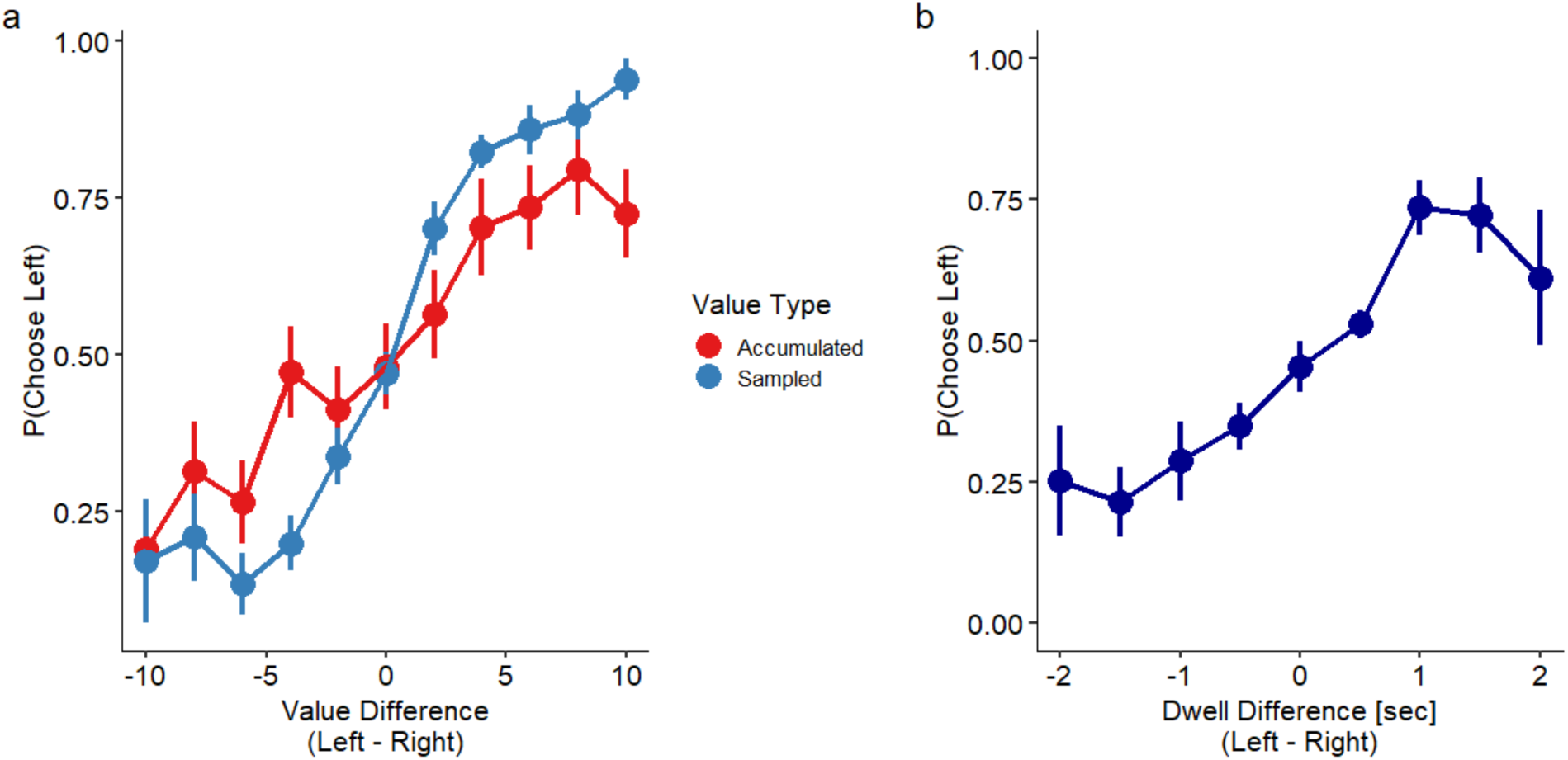
Choice data. (**a**) The probability of choosing left based on Δ*SV* and Δ*AV*. As the value difference becomes greater in favor of one option, the probability of choosing that option increases, for both Δ*SV* and Δ*AV*. (**b**) The effect of gaze on choice. The longer that subjects looked at one lottery over the other, over the course of the whole trial, the more likely they were to choose that lottery.

The larger coefficient for Δ*SV* compared to Δ*AV* is an inevitable consequence of an SSM choice process. In SSMs, a choice occurs when accumulated evidence reaches a threshold. Critically, perceived value for any given sample consists of the true underlying value plus random noise. The final sample (Δ*SV*) is what pushes the accumulated evidence over the threshold, which creates a selection effect: decisions tend to be made when the noise component of Δ*SV* is relatively large and aligned with the ultimate choice, causing the perceived final Δ*SV* to systematically overestimate the true Δ*SV*. As a result, the regression coefficient for the effect of final Δ*SV* on choice is overestimated. In contrast, Δ*AV* represents the sum of all previous evidence, which includes samples that were insufficient to trigger a choice and thus more likely to have noise components that favored the non-chosen option. This means that the perceived Δ*AV* systematically underestimates the true Δ*AV*. As a result, the regression coefficient for the effect of Δ*AV* on choice is underestimated. This creates an inherent asymmetry between Δ*SV* and Δ*AV*: even when the true decision process weights evidence equally over time, regression analyses will show larger coefficients for Δ*SV* than Δ*AV*. For any data generated by an SSM, regressing choice probability on final Δ*SV* and total Δ*AV* would produce a larger coefficient for Δ*SV* due to this threshold-crossing selection effect.

A second feature of SSM data is that easier decisions generally take less time (i.e., fewer samples). Therefore, we expected a negative correlation between the number of samples and the absolute difference in expected value between the two lotteries, as well as a higher probability of terminating a trial when the absolute value difference (|Δ*AV* |) is higher. A mixed-effects regression of log(*n* samples) on the absolute expected value difference between the two lotteries revealed a negative relationship (*β* = -0.025, *SE* = 0.009, *p* = 0.006, two-sided t-test). A mixed-effects logistic regression of P(stop sampling) on |Δ*AV* | also revealed a significant positive relationship (*β* = 0.062, *SE* = 0.009, *p* < 0.001, two-sided t-test). These tests confirm that our subjects were sampling more in more difficult trials.

A third behavioral pattern predicted by the aDDM and other gaze-based SSMs is that individuals should generally choose options that they have looked at more (Cavanagh et al., 2014; Krajbich et al., 2010; Shimojo et al., 2003; Thomas et al., 2019; Westbrook et al., 2020). We thus anticipated a positive correlation between choice and relative dwell time over the course of the whole trial. We added dwell proportion advantage (left – right dwell time divided by total dwell time) to the choice regression and observed a positive effect on choosing the left lottery (*β* =1.642, *SE* = 0.459, *p* < 0.001).

To be certain that gaze-weighted evidence accumulated over the course of the whole trial, and not simply on the final sample of the trial, we excluded the final sample from each trial and re-ran the previous regression. All regressors were significant and positive (Δ*SV*: *β* = 0.290, *SE* = 0.028, *p* < .001; Δ*AV*: *β* = 0.059, *SE* = 0.011, *p* < 0.001; dwell proportion advantage: *β* = 1.636, *SE* = 0.462, *p* < 0.001). Thus, the influence of dwell time on choice occurred over the course of the entire decision, not simply on the final sample (Fig. 3B).

### Neuroimaging results

#### Main analysis: Gaze-based contributions to sampled and accumulated evidence (GLM1)

Our general strategy for the fMRI data was to identify regions with BOLD activity correlating with the time series of |Δ*SV* | or |Δ*AV* |. We primarily focused on the absolute value differences since we were looking for evidence in favor of making any choice, not specifically the left or right choice. This way we could identify the key components of the SSM choice process: the inputs and the integrators. We then tested whether these representations were modulated by gaze.

We tested the following hypotheses: (1) vmPFC and striatum contain input but not integrator representations; (2) the pre-supplementary motor area (pre-SMA), the intraparietal sulci (IPS), and dorsolateral PFC (dlPFC) contain integrator but not input representations; (3) the vmPFC and striatum input representations are modulated by gaze; (4) the pre-SMA, IPS, and dlPFC integrator representations are modulated by accumulated gaze.

To test all four hypotheses, we constructed a general linear model (GLM1) that included both non-gaze-weighted terms (|Δ*SV*| and lagged |Δ*AV* |) and gaze-weighted terms (|Δ*SV_Gaze_*| and lagged |Δ*AV_Gaze_*|). For hypotheses (1) and (2), the variables of interest were absolute sampled value difference (|Δ*SV*|) and absolute accumulated value difference (|Δ*AV*|) where 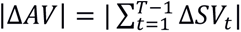, *t* is in units of draws (i.e., pairs of samples), and *T* is the current draw. Note again that we sum |Δ*AV*| to *T-1* in order to exclude the current sample.

For hypotheses (3) and (4), the variables of interest were absolute gaze-weighted sampled values (|Δ*SV_Gaze_*|) and absolute gaze-weighted accumulated values (|Δ*AV_Gaze_*|). The sampled gaze-weighted values (Δ*SV_Gaze_*) were derived by multiplying the proportion of left gaze time with the left value, and the proportion of right gaze time with the right value, within a sample. Accumulated gaze-weighted value differences (Δ*AV_Gaze_*) were the sums of Δ*SV_Gaze_* across samples. Again, |Δ*AV_Gaze_*| did not include the current sample.

These regressors were interacted with boxcar functions covering each sample period (2 seconds). In addition to the regressors of interest, GLM1 contained a stick function for the button press onset, modulated by lagged |Δ*AV*|, as a nuisance regressor event, as well as a boxcar function during the feedback screen, modulated by the value of the received item. We also added motion parameter time series to account for variation due to motion. We report results using FWE-corrected statistical significance of p < 0.05 and a cluster significance threshold of p < 0.005.

### Hypothesis 1: vmPFC and striatum represent sampled but not accumulated value

We first investigated the vmPFC and striatum, regions that we hypothesized represent the inputs (i.e., SV). Looking specifically at BOLD activity in the vmPFC ROI defined in Bartra et al. (2013), we found a positive, but non-significant, correlation with |Δ*SV*| (peak voxel: x = 16, y = 42, z = 12; t = 3.11, p = 0.062) (Fig. 4A, 5A), and unexpectedly, a significant negative correlation with |Δ*AV*| (peak voxel: x = 10, y = 30, z = -4, t = -5.28, p = 0.002). The striatum showed no significant relationship with either |Δ*SV*| or |Δ*AV*| (Fig. 4A, 5A).

**Figure 4.**
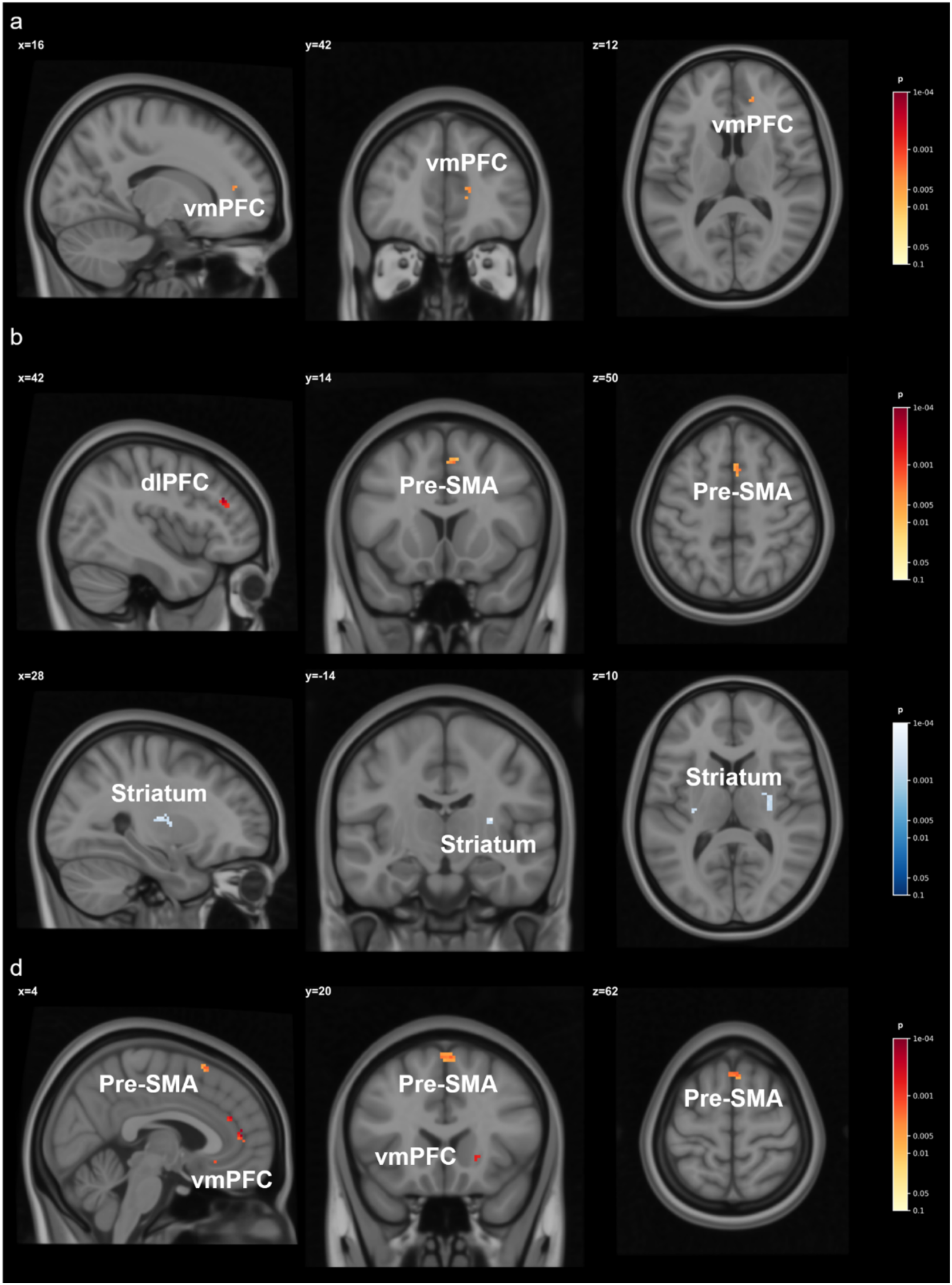
Regions responding to sampled value, accumulated value, and their gaze-weighted variants in GLM1. (a) |Δ*SV*| correlated positively, but not quite significantly, with activity in vmPFC. (b) |Δ*AV*| correlated positively with activity in pre-SMA and dlPFC and negatively with activity in vmPFC (not shown). (c) |Δ*SVGaze*| correlated negatively with activity in the striatum. (d) |Δ*AVGaze|* correlated positively with activity in pre-SMA, vmPFC, and striatum (not shown).

### Hypothesis 2: pre-SMA, IPS, and dlPFC represent accumulated but not sampled value

We next investigated the pre-SMA, IPS, and dlPFC regions, which we hypothesized represent the integrated values (i.e., AV). In the pre-SMA, whose ROI we defined based on Hare et al. (2011), we found a significant, positive relationship between BOLD activity and |Δ*AV*| (peak voxel: x = 6, y = 14, z = 50; t = 3.07, p = 0.023) (Fig 4B, 5A), but no relationship with |Δ*SV*| (Fig. 5A). In the dlPFC, whose ROI we defined based on Hare et al. (2011), we found the same pattern as the pre-SMA: there was a significantly positive relationship between BOLD activity and |Δ*AV*| (peak voxel: x = 42, y = 34, z = 28; t = 4.29, p = 0.020) (Fig. 4B, 5A), and no relationship between BOLD activity and |Δ*SV*| (Fig. 5A). In the IPS, identified with the Harvard-Oxford Cortical Structural Atlas, we found no relationship with |Δ*AV*| or |Δ*SV*| (Fig. 4B, 5A).

**Figure 5.**
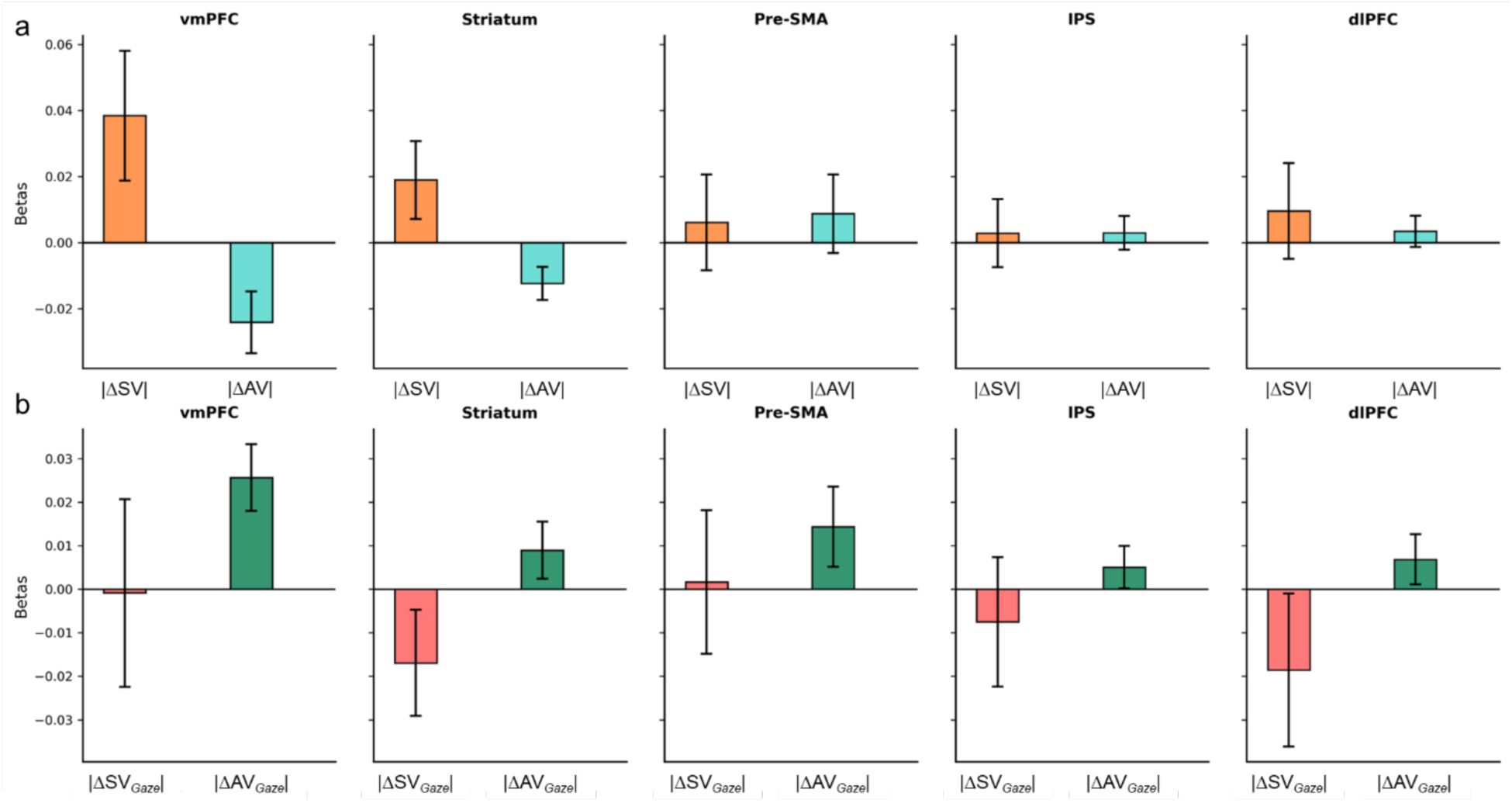
GLM1 beta plots from the vmPFC, striatum, pre-SMA, IPS, and dlPFC. Displayed are regression coefficients from each region for (a) non-gaze-weighted signals: absolute sampled value difference (|Δ*SV*|) and absolute lagged accumulated value difference (|Δ*AV*|), and (b) gaze-weighted signals: gaze-weighted sampled value (|Δ*SVGaze*|) and absolute lagged gaze-weighted accumulated value (|Δ*AVGaze|*). Note: Bar heights reflect mean beta estimates averaged across all voxels within each ROI. Statistical significance was determined using permutation tests with FWE correction, which identify spatially localized, reliable effects within each ROI.

In summary, these gaze-free regressors yielded only weak support for the expected vmPFC result, no support for the expected striatum and IPS results, and strong support for the hypothesis that pre-SMA and dlPFC represent integrators but not inputs.

### Hypothesis 3: vmPFC and striatum represent gaze-modulated sampled values

We next asked whether the activity in these regions was affected by gaze. The aDDM (and other gaze-weighted SSMs) predict that gaze to one option should amplify that option’s value relative to the other option (Krajbich et al., 2010; Thomas et al., 2019; Westbrook et al., 2020). Consider a simple model where an option’s value is weighted by the proportion of time during which it is looked at. Imagine two trials with the same pair of values, 7 on the left and 3 on the right. In Trial A, the subject looks left 30% of the time and right 70% of the time. In Trial B, the subject looks left 70% of the time and right 30% of the time. In Trial A, the net input value (“drift rate”) would be |0.3·7-0.7·3|=0. In Trial B, the drift rate would be |0.7·7-0.3·3|=4. In Trial A, the value advantage for the left option is canceled out by the gaze advantage for the right option. In Trial B, both the value and gaze advantage favor the left option, leading to strong evidence in favor of a left choice. In sum, there is stronger evidence when gaze difference is aligned with value difference. This should be true for both Δ*SV* and Δ*AV*, though Δ*SV* is only affected by gaze during the current draw, while Δ*AV* is affected by gaze over the entire trial.

In the vmPFC, we found no significant correlation with |Δ*SVGaze*|, but a positive correlation with |Δ*AVGaze*| (peak voxel: x = 6, y = 50, z = 12; t = 5.52, p = 0.002) (Fig. 4C, 5B).

Surprisingly, the striatum showed a significant negative correlation with |Δ*SVGaze*| (peak voxel: x = 30, y = -14, z = -12; t = -4.88, p = 0.014) (Fig. 4C, 5B), and a positive correlation for |Δ*AVGaze*| (peak voxel: = -4, y = 6, z = -10; t = 4.22, p = 0.047) (Fig. 5B). These results are notable because the striatum did not show sensitivity to non-gaze-weighted signals, suggesting that gaze modulation reveals striatal involvement in value processing that would otherwise be obscured.

### Hypothesis 4: pre-SMA, IPS, and dlPFC accumulated value representations are modulated by gaze

In the pre-SMA we found a significantly positive relationship with |Δ*AVGaze*| (peak voxel: x = 4, y = 20, z = 62; t = 3.34, p = 0.033) (Fig. 4D, 5B). The IPS and dlPFC, on the other hand, did not show significant correlations with |Δ*AVGaze*| nor with |Δ*SVGaze*| (Fig. 5B).

In summary, these gaze-modulated regressors indicate that the vmPFC, striatum, and pre-SMA all appear to represent accumulated value, while the IPS and dlPFC show no such effects.

### Gaze allocation and value alignment (GLM2)

To more directly test whether accumulated evidence signals were modulated by accumulated gaze allocation throughout a trial, we conducted additional, exploratory analyses. Specifically, we ran a GLM that incorporated the following two terms: accumulated dwell advantage and Δ*AV* × accumulated dwell advantage, in addition to Δ*SV*, the current gaze location, and Δ*SV* × current gaze location.

We calculated accumulated dwell advantage as follows: For each sample *t*, accumulated dwell advantage is the cumulative difference in gaze allocation up to sample *t*-1, calculated as (total dwell left – total dwell right) / (total dwell left + total dwell right). This is a continuous measure from -1 (all previous gaze to right) to +1 (all previous gaze to left).

We also included the interaction between accumulated dwell advantage and Δ*AV* (i.e., signed accumulated evidence). This interaction term is positive when gaze is primarily to the left and left has more value or when gaze is primarily to the right and right has more value. This interaction term directly tests whether brain regions encoding accumulated evidence are modulated by the history of gaze allocation. This approach allows us to examine the interaction effect more explicitly as a separate regressor rather than having it embedded within the value calculation itself.

This GLM revealed a positive correlation between pre-SMA activity and the Δ*AV* × accumulated dwell advantage term (peak voxel: x = 8, y = 10, z = 58; t = 3.01, *p* = 0.028). The striatum also showed a correlation with this term (peak voxel: x = -16, y = 10, z = -6; t = 4.07, *p* = 0.016). Additionally, activity in the dlPFC was positively correlated with Δ*SV* (peak voxel: x = -36, y = 34, z = 22; t = 3.96, *p* = 0.016), not modulated by dwell advantage. No other ROIs showed significant relations.

This analysis provides additional evidence that the pre-SMA and striatum encode accumulated value signals that are modulated by the history of gaze allocation.

### Accounting for individual difference in attentional discounting (GLM3)

In our first set of analyses, we implicitly assumed complete discounting of non-fixated information, in contrast with previous studies that have generally found only partial discounting (Krajbich et al., 2010; Sepulveda et al., 2020; Smith & Krajbich, 2019; Westbrook et al., 2020). To verify that our results are robust to inter-subject variability in attentional discounting, we estimated subject-level attentional discounting parameters and then re-estimated our original GLM with new, recalculated gaze-weighted value regressors.

Following Smith, Krajbich, and Webb (2019), for each individual sample within a trial, we computed:

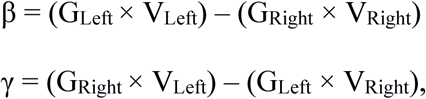

where GLeft and GRight represent the proportion of time spent gazing left versus right within that specific sample, and VLeft and VRight are the instantaneous values of the left and right options. We then averaged these sample-level β and γ values across all samples within each trial to obtain trial-level regressors. We then ran a mixed-effects logistic regression predicting choice (left vs. right) as a function of β and γ and then calculated subject-specific values of θ = γ/β. Across our sample (N=20), we found mean θ = 0.77 (SD = 0.21, range = 0.55–1.25).

Next, for the GLM, we computed θ-weighted sampled-value (|Δ*SV_θ_*|) as:

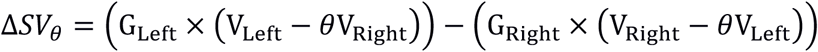

Similar to the original GLM, we computed an accumulated value signal, |Δ*AV_θ_*|, based on the lagged sum of previous samples’ |Δ*SV_θ_*|.

We found a marginally positive effect of |Δ*SV_θ_*| in the vmPFC (peak voxel: x = -14, y = 44, z = -12; t = 3.30, *p* = 0.052), a positive but not significant effect in the IPS (peak voxel: x = - 34, y = -28, z = 34; t = 3.82, *p* = 0.068), and no significant effects in other ROIs.

In contrast, we found significant positive relationships between |Δ*AV_θ_*| and activity in the pre-SMA (peak voxel: x = 4, y = 12, z = 50; t = 6.13, *p* < 0.001), dlPFC (peak voxel: x = 40, y = 34, z = 24; t = 6.70, *p* < 0.001), and IPS (peak voxel: x = 40, y = -50, z = 40; t = 5.55, *p* = 0.004). Notably, we also observed both positive and negative, but not significant, relationships between |Δ*AV_θ_*| and activity in the vmPFC (positive peak voxel: x = 6, y = 38, z = 26; t = 3.30, *p* = 0.059; negative peak voxel: x = 10, y = 30, z = -4; t = -3.06, p = 0.080). No other notable contrasts emerged.

In summary, these analyses provide some evidence that the vmPFC encodes gaze-weighted sampled and accumulated value signals, while the pre-SMA, dlPFC, and IPS encode gaze-weighted accumulated value signals.

### Modeling a non-uniform temporal weighting function (GLM4)

While sequential sampling models like the DDM assume equal weighting of information during evidence accumulation, other models allow for information sampled at different time points to differentially impact choice. For example, information that arrives early (i.e., primacy) or late (i.e., recency) can preferentially influence decision-making (Usher & McClelland, 2001). This could especially be the case in our task where information has to be integrated over a long period of time.

To account for temporal biases during evidence accumulation, we fit participant data with a model that incorporates primacy and recency biases into the sequential sampling process (Galdo et al., 2022; Pooley et al., 2011). Within a trial *t,* the weight of sample *i* is determined by the following temporal weighting function:

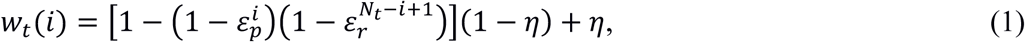

where *ε_p_* and *ε_r_* are weights on primacy and recency, respectively, *η* is a lower bound on the weight of any sample, and *N_t_* is the number of samples on trial *t*.

In the context of the current experiment, we assume that decision-makers accumulate evidence (Δ*AV*) based on the sum of sampled evidence (Δ*SV*) weighted by the temporal weighting function:

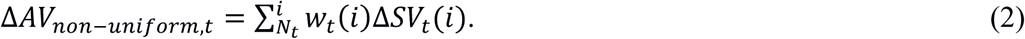

At the time of choice, the decision-maker chooses according to the logit function:

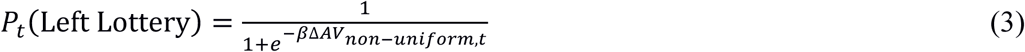

with inverse-temperature parameter *β* governing the how strongly Δ*AV_non_*_−*uniform*,*t*_ governs the selection of the higher value lottery.

For models incorporating visual attention, the only modification is to use Δ*SVGaze* instead of Δ*SV*.

### Behavioral model results

We observed a substantial recency bias, with all participants showing *ε_r_* > *ε_p_* when using Δ*SV* and all but one participant showing *ε_r_* > *ε_p_* when using Δ*SVGaze* (Fig. 6) as inputs. Independent samples t-tests (with the Welch approximation to degrees of freedom) confirmed that participants’ *ε_r_* parameters were significantly larger than their *ε_p_* parameters, for both Δ*SV* (*t*(36) = 12.54, *p* = 10^−14^) and Δ*SVGaze* (*t*(20) = 7.25, *p* = 10^−6^) inputs. Goodness-of-fit measures based on the Bayesian Information Criterion (BIC) also favored a non-uniform temporal weighting function (No gaze: BIC = 1262; Gaze: BIC = 1593) over a uniform temporal weighting function (No gaze: BIC = 1307; Gaze: BIC = 1658). However, it is worth noting that the recency bias is surely over-estimated based on our parameter-recovery exercise (Methods) and the bias introduced by allowing participants to choose when to stop collecting evidence (Discussion).

**Figure 6.**
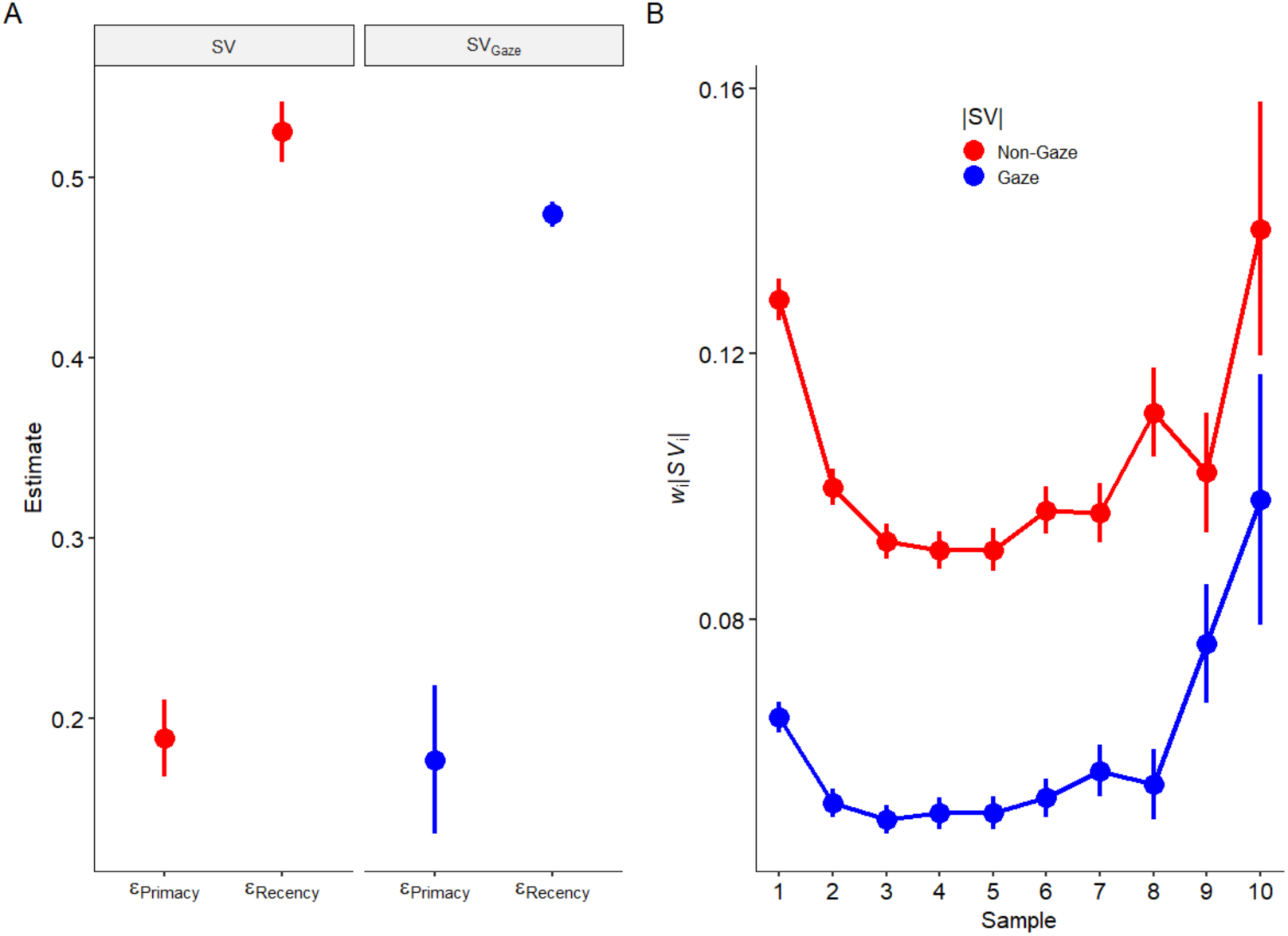
Non-uniform temporal weighting. In both gaze-weighted and non-gaze-weighted models, participants showed stronger recency than primacy effects, both in terms of **(A)** the model parameters, and **(B)** the resulting temporal weighting functions averaged across all trials. Error bars are standard errors clustered by participant.

### fMRI model results

To identify brain regions that encoded |Δ*AVnon-uniform|*, we tested whether activation parametrically varied as a function of these new values using updated versions of GLM1 in the previously defined ROIs. Within each ROI, we used a FWE corrected threshold of *p* < 0.05 and cluster-forming threshold of *p* < 0.005 with 5000 permutations.

Here we again found a positive, but not quite significant, correlation between |Δ*SV*| and activity in the vmPFC (peak voxel: x = -14, y = 42, z = -12; t = 2.95, *p* = 0.088). We also observed a surprising negative correlation between |Δ*SVGaze*| and activity in the striatum (peak voxel: x = 30, y = -14, z = 12; t = -4.61, *p* = 0.027).

Turning to non-gaze-weighted accumulated value, we found a significant positive correlation between |Δ*AV*non-uniform| and BOLD activity in the IPS (peak voxel: x = 46, y = -54, z = 44; t = 4.35, *p* = 0.032). Unexpectedly, we also observed a marginally negative correlation in the striatum (peak voxel: x = 6, y = 4, z = -10; t = -3.79, *p* = 0.056).

For gaze-weighted |Δ*AV*non-uniform|, we found significant correlations with BOLD activity in the pre-SMA (peak voxel: x = 6, y = 18, z = 52; t = 4.21 *p* = 0.006), dlPFC (peak voxel: x = 38, y = 30, z = 26; t = 4.28, *p* = 0.003), and vmPFC (peak voxel: x = 8, y = 38, z = 24; t = 3.62, p = 0.032).

In summary, these analyses provide additional evidence that the pre-SMA, dlPFC, and vmPFC encode gaze-weighted accumulated value signals.

## Discussion

In this article we presented results from a simultaneous eye-tracking and fMRI study of value-based decision-making, using an expanded-judgment task where subjects sampled from, and then chose between food lotteries. Across each of our analyses, we consistently found that the pre-SMA represents accumulated value signals that are modulated by gaze. We also observed some evidence for gaze-weighted accumulated value signals in dlPFC, vmPFC, and the striatum, while activity in the IPS correlated only with gaze-free accumulated values. Activity correlating with sample value was weak and inconsistent, occurring primarily in the vmPFC and (negatively) in the striatum, in contrast to prior findings in these regions (Lim et al., 2011; McGinty et al., 2016).

These results provide novel evidence for the neural mechanisms underlying the SSM process, as exemplified by the DDM, which appears to govern many types of decisions (Busemeyer, 1985; Ratcliff, 1978). The gaze modulation of accumulated value signals in the pre-SMA and perhaps other regions, provides critical evidence that these regions indeed represent accumulated evidence, as opposed to unchosen values (Boorman et al., 2011; Kolling et al., 2016; Wittmann et al., 2016), decision conflict (Frank et al., 2015; Frömer et al., 2019; Kaanders et al., 2021; Shenhav et al., 2014, 2016; Vassena et al., 2020), or time on task (Grinband et al., 2011; Holroyd et al., 2018; Mumford et al., 2024). While accumulated evidence is typically correlated with these other measures, we were able to dissociate them by taking advantage of the fact that accumulated evidence, but not the other measures, are modulated by gaze location.

Value is amplified by gaze (Smith & Krajbich, 2019; Westbrook et al., 2020), leading to stronger value signals in the brain when the decision maker is looking at the higher value option.

Conflict-based models make qualitatively different predictions. Regions implementing conflict monitoring should show increased activity when options are similar in value, regardless of time. The conflict account predicts that BOLD activity should scale with inverse value difference: smaller |ΔV| → higher conflict → higher BOLD (Shenhav et al., 2014, 2016). In simple choice tasks, high conflict and high accumulated value are both associated with long response times (Pisauro et al., 2017), leading to ambiguity about how to interpret purported neural correlates of accumulated value. In our task we avoided this ambiguity by analyzing the effect of accumulated value at each point in time, not just at the moment of decision. With this approach, conflict should be inversely correlated with accumulated value (as higher accumulated evidence indicates less similarity between options). Moreover, the conflict account makes no predictions about how BOLD activity should be modulated by gaze allocation for a given set of option values.

A more serious concern is the potential confound with time-on-task BOLD activity. Accumulated value inevitably increases with time within a trial, leading to a correlation between the two variables (Grinband et al., 2011; Holroyd et al., 2018; Mumford et al., 2024). This is where the gaze data were particularly important. Time-on-task regions should show no relation with gaze allocation patterns. After accounting for non-gaze-weighted accumulated value, only accumulator regions, and not time-on-task regions, should show a relationship with gaze-weighted accumulated value. The results of our analyses provide exactly such evidence: pre-SMA activity, and to some extent striatal activity, was positively correlated with gaze-weighted accumulated value, even when accounting for previous gaze history and individual differences in attention discounting.

Representation of unchosen values (Boorman et al., 2011; Kolling et al., 2016; Wittmann et al., 2016) might explain some of the unexpected negative correlations that we sometimes observed between our value regressors and activity in the vmPFC, striatum, and IPS. One potential explanation for these negative results is the fact that when value difference is large, one of the two values is often very low. In other words, there is a negative correlation between |Δ*V*| and min {*V*_1_, *V*_2_}. If some neurons in these regions represent the values of individual items or lotteries (Padoa-Schioppa & Assad, 2006; Rich & Wallis, 2016), then we might expect to see some activity negatively correlated with |Δ*V*|. This activity would align with a representation of the unchosen values.

Our findings were made possible by considering the role that visual attention plays in the decision process. While SSMs capture choice behavior and RTs extremely well, most do not consider the effects of attention. Attention is thought to shift over the course of the decision, amplifying the attended inputs and/or inhibiting the non-attended inputs (Diederich, 1997; Johnson & Busemeyer, 2005; Roe et al., 2001). These shifts in attention are reflected in eye-movements (Hoffman & Subramaniam, 1995), which affect choice outcomes (Fiedler et al., 2013; Fiedler & Glöckner, 2015; Folke et al., 2016; Glaholt & Reingold, 2009; Gwinn et al., 2019; Janiszewski et al., 2013; Jiang et al., 2016; Kim et al., 2012; Konovalov & Krajbich, 2016; Lopez-Persem et al., 2016; Orquin & Mueller Loose, 2013; Pärnamets et al., 2015; Polonio et al., 2015; Russo & Leclerc, 1994; Sheng et al., 2020; Shi et al., 2012; Smith & Krajbich, 2018; Stewart et al., 2016; Tavares et al., 2017; Towal et al., 2013; Vaidya & Fellows, 2015; Vanunu et al., 2021; Venkatraman et al., 2014; J. T. Wang et al., 2010), and their effect on the choice process is captured by the attentional drift diffusion model (aDDM) and other related SSMs (Ashby et al., 2016; Fisher, 2021; Glickman et al., 2019; Jang et al., 2021; Krajbich et al., 2010; Li & Ma, 2021; Smith & Krajbich, 2019; Teoh et al., 2020; Westbrook et al., 2020; Yang & Krajbich, 2023; Zilker & Pachur, 2023).

Neural implementations of SSMs generally require at least two sets of neurons, one set to represent the current information from the stimuli and a second set to integrate that information over time. In the current study, the information being used to make decisions was subjective value (i.e., utility). A large body of work has implicated the vmPFC and striatum in representing value (Bartra et al., 2013; Boorman et al., 2009; Chib et al., 2009; Levy & Glimcher, 2011; Schoenbaum et al., 1998). Our results partially confirm that the vmPFC represents value information. The integrated value information, which is what ultimately determines the decision, is primarily encoded in the pre-SMA, though the vmPFC, striatum, and dlPFC may also be involved. The results in the vmPFC are consistent with work by some researchers, arguing that in addition to representing value, the vmPFC also represents the decision variable (Padoa-Schioppa & Assad, 2006; Strait et al., 2014). The results in the dlPFC are also consistent with research in perceptual and value-based decision making, where research has argued that the dlPFC is involved in accumulation and comparison processes (Busemeyer et al., 2019; Gluth et al., 2012; Heekeren et al., 2006; Hutcherson & Tusche, 2022). Future work will need to further disentangle the role that these regions play in the choice process.

By using eye-tracking, our study extends previous work connecting computational models to fMRI data. Our results partially align with Hare et al.’s (2011) proposed neural model in which the vmPFC provides inputs to the pre-SMA and IPS, with the pre-SMA also exchanging information with the dlPFC. While the Hare model was based on dynamic causal modeling results, our task provides a more direct test of the proposed neural network.

Additionally, our eye-tracking data allows us to identify additional features of the network. We find that the pre-SMA and dlPFC are sensitive to gaze-weighted accumulated value, while the IPS is not. The reason for this distinction between the frontal regions and IPS is unclear, but it does suggest that the pre-SMA is more likely to be the final decision-making region, consistent with some recent studies (Juechems et al., 2017; Pisauro et al., 2017; Rodriguez et al., 2015; Rouault et al., 2019; Aquino et al., 2023). Of course, the recruitment of the pre-SMA may be because our subjects made their decisions with a button press, which is supported by the correlations between accumulated value and motor cortex activity. Had our study required eye-movements to indicate a choice, we may very well have observed integrator activity in other regions such as the frontal eye fields or posterior parietal cortex (O’Connell et al., 2018). However, the activity in dlPFC may be independent of response modality, consistent with findings in perceptual decision making (Heekeren et al., 2006).

A major advantage of this study is its use of a task designed to slow down the decision process and force sequential integration of information. Such expanded judgment tasks have been used to study SSM assumptions in perceptual decision-making, more recently in combination with neural recordings, but mostly with electrophysiology in rats and monkeys (Brunton et al., 2013; Cisek et al., 2009; Tsetsos et al., 2012; T. Yang & Shadlen, 2007).

On the other hand, one concern with longer decision times is that decision-makers might either under-weight (i.e., forget), or put too much weight on, early information. Our analysis of subjects’ temporal weighting functions did reveal a primacy effect, where the first sample in each trial was overweighted, as well as a recency effect, where the last samples in each trial were also overweighted. Nonetheless, using a temporal weighting function with these primacy and recency effects did not substantially change the conclusions from our fMRI analysis. While these analyses did reveal weak support for accumulator dynamics in the vmPFC, these results should be interpreted with caution because adding recency effects into the accumulated-value signal increased the correlation between |Δ*SV*| and |Δ*AV*| from 0.12 to 0.21. Moreover, the recency effect is surely overestimated due to well-known statistical artifacts (Mullett & Stewart, 2016).

In short, because decision-makers tend to terminate the decision process after strong pieces of evidence, and because a strong piece of evidence will tend to have a large noise component, information that appears at the end of a trial will appear to have a stronger influence on choices than it should.

Another potential limitation of our study was the number of participants. While our sample size of 20 subjects is modest by current neuroimaging standards, the within-subject statistical power from our extended decision paradigm (∼380 observations per subject), combined with hypothesis-driven ROI analyses and multiple comparisons correction, provides confidence in our core findings. Nevertheless, replication with larger samples would be valuable, particularly for more fully characterizing null effects and marginal findings.

Taken together, we separated sampled input values from the overall decision value, or accumulated value, and found a network of brain regions that are involved in an aDDM-like choice process. This process involves passing sampled input values to an integrator which responds to not only the values themselves, but also to the gaze-modulated values. These results indicate that gaze effects on value representations are not epiphenomenal; rather, they reflect how gaze is incorporated into the decision process, affecting how we perceive the value of each option over the course of the entire decision.

## Materials and Methods

### Experimental Design and Statistical Analyses

#### Subjects

Twenty-eight undergraduate students at The Ohio State University participated in this study. Due to time constraints, 4 subjects were unable to finish all 3 runs of the scan. One subject was excluded for not choosing in line with their ratings. This left 23 subjects in the analysis (14 men, 9 women, average age: 22.61). Another 3 subjects could not be calibrated on the eye-tracker and so are discarded from any analyses involving eye tracking (leaving 13 men, 7 women, average age: 22.9). All subjects were right-handed, had normal or corrected-to-normal vision, and no history of neurological disorders. This study was approved by The Ohio State Biomedical Sciences IRB.

#### Stimuli and Tasks

*Rating Task*: Outside of the scanner, subjects rated 148 food items using a continuous rating scale from -10 (extreme dislike) to 10 (extreme like), with 0 being indifference towards an item. To choose their preferred rating, subjects moved the mouse across the rating scale and then clicked the left mouse button when the cursor was at their desired value for the item. We used an incentivization procedure for these ratings. There was a 50% chance that the rating task would be used to determine the subject’s reward. In such cases, the computer would randomly select two foods and the subject would receive the one with the higher rating. If both items were rated negatively, the subject would not receive any food. Negative values (-10 to 0) were excluded from the choice task except for two subjects who did not have enough positively valued items.

#### *Choice Task:* Once in the MRI scanner, subjects chose between pairs of lotteries

Lotteries were constructed by creating 1000 potential lotteries of randomly selected items evenly split between having 3, 4, 5, or 6 items. Each item in a lottery was then assigned a probability of being drawn, with probabilities summing up to 1 in each lottery. For each of these lotteries we calculated its expected utility by multiplying the subjective value (i.e., rating) of each item (*V_i_*) in the lottery by its associated probability (*P_i_*) of being drawn, and summing the results

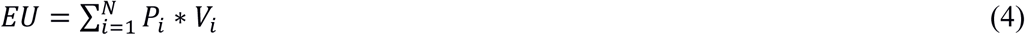

where N is the total number of items in that lottery. Subjects were not told these probabilities, nor were they told which items were in each lottery – they had to learn this every trial.

We tracked subjects’ eye movements in the scanner using an Eyelink 1000 plus (SR Research) set at 500 Hz. Eye position was monitored with the camera and infrared source reflected in the mirror attached to the head coil. The eye tracker was calibrated at the beginning of the session.

Food items were sampled from the lotteries and presented on the screen one pair at a time. Each draw was presented for 2 seconds, followed by a fixation cross for 2-6 seconds. This process repeated until the subject made a choice. Once a subject was ready to choose a lottery, they used the index finger of their left (right) hand to press a button corresponding to the left (right) lottery. They were then presented with a random item from the lottery they had chosen (on the same side of the screen as the chosen basket) for 2 seconds, indicating the food they would receive from this trial, should it be randomly selected at the end of the study (Fig. 1).

We constructed each trial’s sequence of items pseudo-randomly to minimize the correlation between the sampled value signal (|Δ*SV*|; i.e., the absolute difference in sampled input values) and the accumulated value signal (|Δ*AV*|; i.e., the absolute difference in accumulated values). For the first draw in each trial, sampled and accumulated value signals are equal. On subsequent draws, the |Δ*SV|* diverges from the |Δ*AV|* signal, yielding two distinct time courses to look for in the fMRI data (Fig. 2). Across subjects, |Δ*SV*| and |Δ*AV*| had an average correlation of 0.31 (SD = 0.10, min = 0.08, max = 0.45), while |Δ*SV*| and lagged |Δ*AV*| (i.e., the variables in our GLMs) had an average correlation of 0.07 (SD = 0.08, min = -0.09, max = 0.17). See Figure 7 for the full correlation matrix of value signal regressors calculated across subjects and samples.

**Figure 7.**
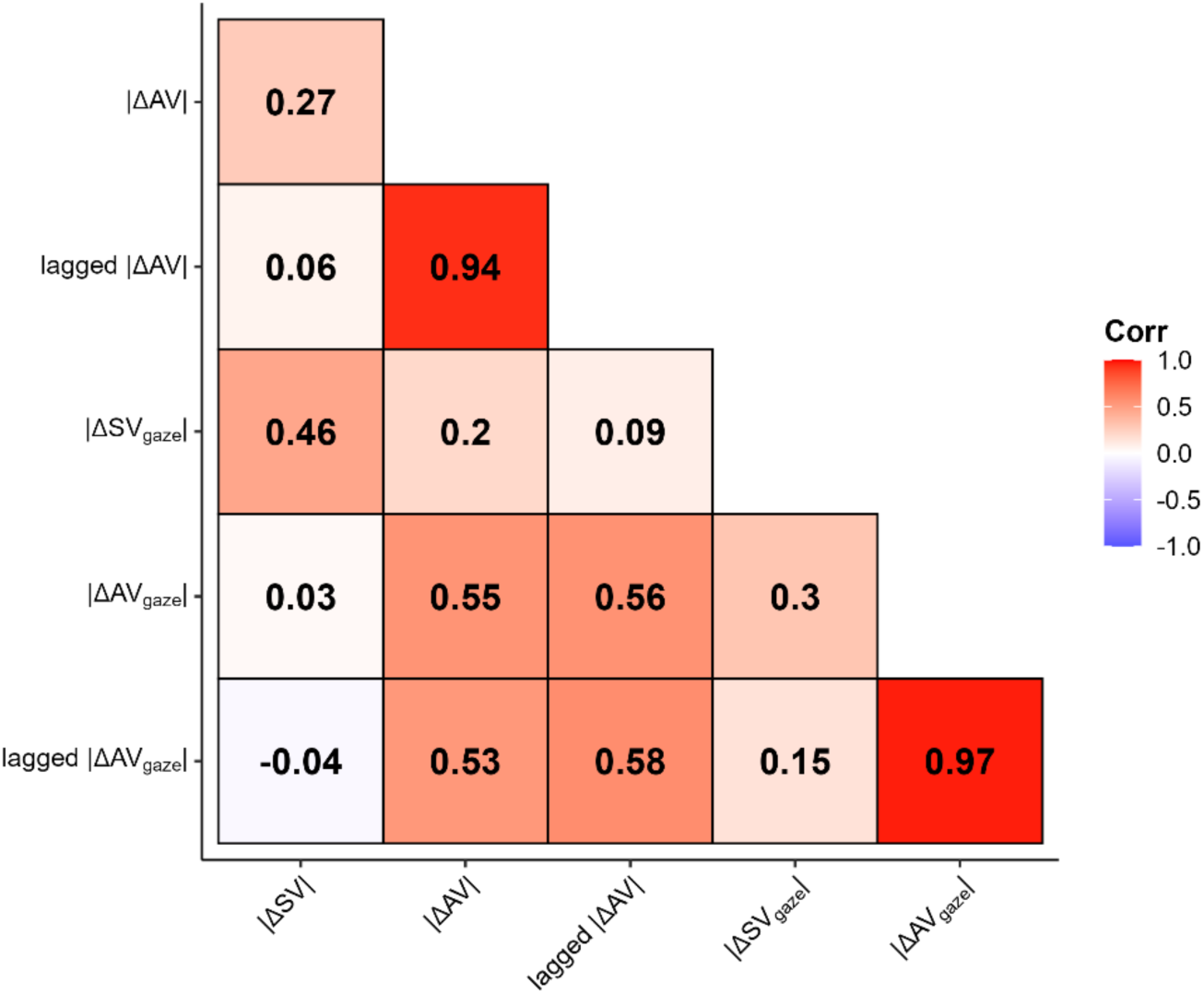
Sample-level correlations between value signal regressors. Pearson correlations computed across samples for different quantifications of sampled value (Δ*SV*) and accumulated value (Δ*AV*). Values represent correlations between absolute magnitudes of each measure. |Δ*SV|* = absolute sampled value difference; |Δ*AV|* = absolute accumulated value difference; lagged |Δ*AV|* = absolute accumulated value difference up to the previous sample; |Δ*SVGaze|* = gaze-weighted sampled value; |Δ*AVGaze|* = gaze-weighted accumulated value; lagged |Δ*AVGaze|* = gaze-weighted accumulated value from the previous sample.

The task structure was designed to incentivize subjects to average 7 samples per trial. They had 45 minutes to make 60 choices, and any trials that they did not complete by the end of the experiment were made for them randomly by the computer. Any trials they completed beyond 60 were simply added to the pool of potentially rewarded trials. At the end of the study, there was a 50% chance that one of the choice trials would be randomly selected for payment (otherwise the rating task was selected for payment), in which case the subject received the corresponding food from that trial.

Outside of the scanner, subjects first completed a 5-minute practice section where they chose between baskets made up of cars. After each trial, they were given feedback on how many samples they took and were reminded that the goal was to take 7 samples on average. These choices were not incentivized.

#### Temporal-weighting-function model fits

We estimated model parameters using an iterative maximum a posteriori (MAP) approach (Huys et al., 2011; Wittmann et al., 2020). This method improves upon maximum likelihood estimation (MLE) by simultaneously estimating parameters at both the subject- and group-level. This hierarchical procedure constrains subject-level parameters and reduces the influence of outlier data.

Group-level parameters were initialized with uninformative Gaussian priors with mean of 0.1 and variance of 100. For all models, *η* was held constant at 1. During the expectation step, we estimated model parameters (*ε_p_*, *ε_r_*, *β*) for each participant using MLE and calculated the log-likelihood of their choices given the model parameters. During the maximization step, we calculated the maximum posterior probability based on the observed choices and prior group-level parameters, and then updated the group-level parameters to generate posterior parameter estimates. These posterior parameter estimates were then used as the priors in subsequent steps in this procedure. We iteratively repeated the expectation and maximization steps until convergence of the posterior likelihood summed over group-level parameters exceeded a change of less than 0.0001 from the previous iteration (for a maximum of 800 iterations). During this procedure, bounded free parameters were transformed from Gaussian space to native model space using link functions (e.g., sigmoidal function for *ε_p_*, *ε_r_*) to ensure accurate estimation near the bounds.

We assessed parameter recovery of this model on simulated choices using best-fitting model parameters. We then refit the simulated choices using the same MAP process described above. We found strong Pearson’s *r* correlations between the generated and estimated parameter values (*r* > .8; Fig. 8), though both parameters were systematically over-estimated.

**Figure 8.**
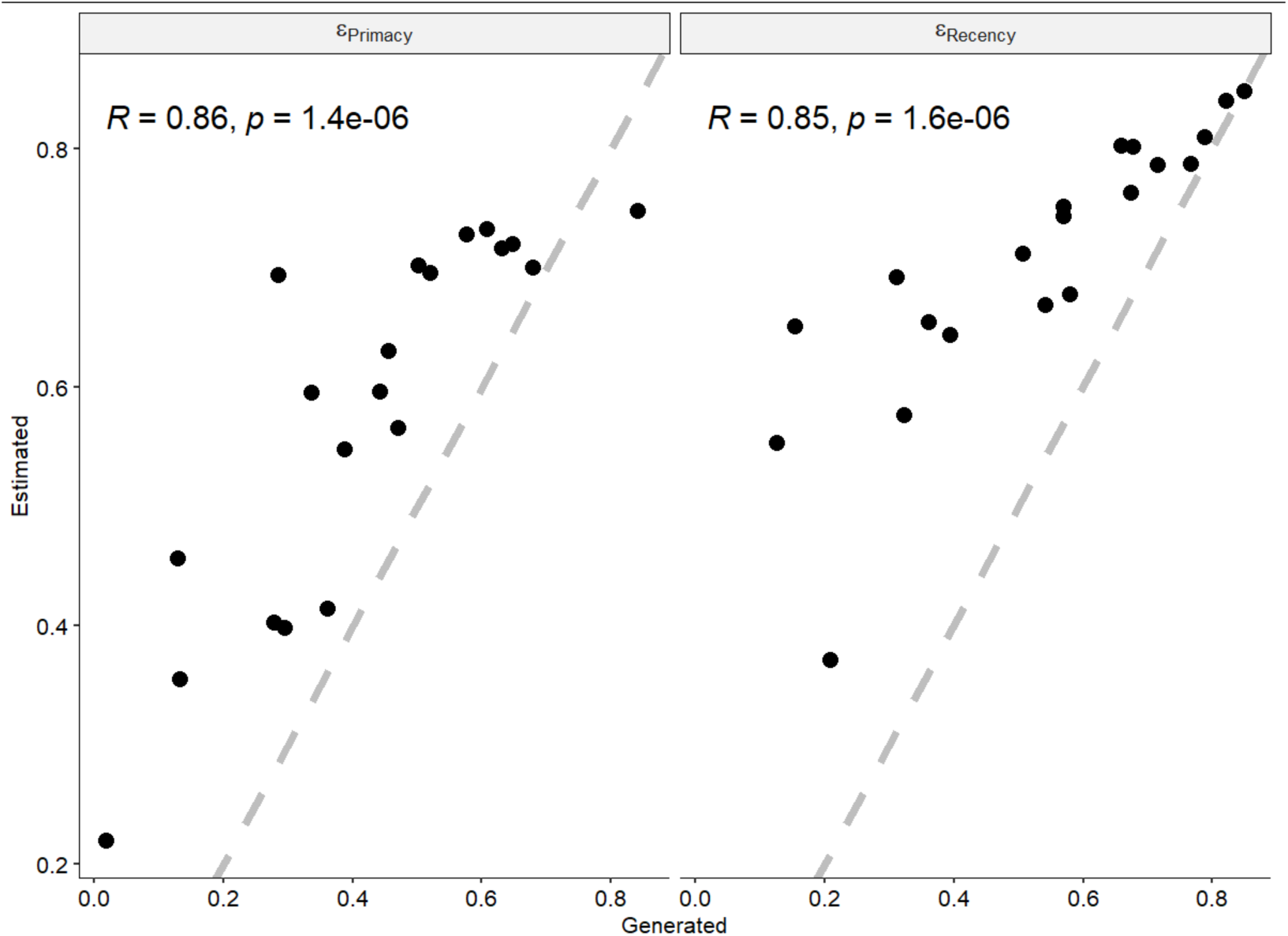
Correlation plots illustrating parameter recovery.

#### MRI Data Acquisition

MRI scanning was carried out at the OSU Center for Cognitive and Behavioral Brain Imaging. We used a 3T Siemens Magnetom Prisma scanner with a 32-channel head array coil to collect the neural data. Functional data were acquired with a T2*-weighted gradient-echo sequence (48 slices, interleaved, with a field of view of 1554x1554, with an in-plane resolution of 3 mm isotropic and a 3mm slice gap, TR = 2600 ms, TE = 28 ms, 80° flip angle). Slices were oriented such that the anterior side of the acquisition was raised dorsally by 30 degrees compared to the line formed by joining the anterior commissure to the posterior commissure. A high-resolution MPRAGE anatomical scan (256 slices, field of view 224x256, with an in-plane resolution of 1 mm and no slice gap, TR = 1900 ms, TE = 4.44 ms, 12° flip angle) was also acquired for each participant. Each participant was scanned in one 1.5-hour session, which included the three experimental runs (15 minutes each) and the high-resolution MPRAGE anatomical scan. Additionally, a resting state scan (5 minutes) and a DTI scan were acquired, but these data are not presented here. Stimuli were presented using Psychtoolbox (Brainard, 1997; Kleiner et al., 2007; Pelli, 1997) for MATLAB (MATLAB, 2016), and displayed with a DLP projector onto a screen mounted in the rear of the scanner bore.

#### MRI Preprocessing and analyses

Statistical parametric mapping (SPM12, Update Rev. Nr. 6905; Functional Imaging Laboratory, University College London) was used to carry out the preprocessing of fMRI data. First, we corrected for the different slice times per echo planar image (EPI) across the total volume (using the bottom slice as a reference) and then realigned each volume in a run to the mean EPI volume from that run. Next, the anatomical scan was coregistered with the MNI average of 152 brains template, and the mean EPI per run was used to coregister all functional scans to this coregeistered anatomical scan. In order to warp the EPIs to MNI space, SPM12’s *normalise* function was applied to the coregistered anatomical scan and the resulting warping parameters were applied to the coregistered EPIs. The resulting images were smoothed using an isometric Gaussian kernel (8 mm full width at half maximum). First level GLMs were run using SPM on each subject individually, including contrasts of interest. Runs were combined using *fslmerge*. We then used the Statistical non-Parametric Mapping toolbox (e.g., SnPM013.1.09) function to run second-level, non-parametric significance tests with threshold-free clustering and family wise error (FWE) correction to find significant clusters for the described effects (Nichols & Holmes, 2002). We report the results of 5000 permutations using FWE-corrected statistical significance of *p* < 0.05 and a cluster significance threshold of *p <* 0.005.

We used a canonical hemodynamic response function (HRF) without time derivatives. We modeled the noise as an AR(1) process. We additionally used a high-pass filter set to 128 s. We also used global signal normalization with a value of 0.8. We used no corrections for susceptibility distortions. All general linear models (GLM) included variants of |Δ*SV*| and lagged |Δ*AV*|, either gaze weighted or not, interacted with boxcar functions covering each sample period (2 seconds) – see details below. In addition to the regressors of interest, each GLM contained the following nuisance terms: a stick function for trial number, a stick function for the button press onset modulated by lagged |Δ*AVGaze*|, and a boxcar function during the feedback screen, modulated by the value of the received item. We also added motion parameter time series to account for variation due to motion.

In our primary analyses (GLM1), in addition to the nuisance terms, we included the following regressors: |Δ*SV*|, lagged |Δ*AV*|, |Δ*SVGaze*| and lagged |Δ*AVGaze*|.

In our analyses of gaze allocation history (GLM2), in addition to the nuisance terms, we included the following regressors: Δ*SV*, lagged Δ*AV*, current gaze location, accumulated dwell advantage, Δ*SV* × current gaze location, and lagged Δ*AV* × accumulated dwell advantage.

In our analysis incorporating individual differences in attentional discount (GLM3), in addition to the nuisance terms, we included the following regressors: *θ*-weighted |Δ*SV*| and lagged *θ*-weighted |Δ*AV*|.

In our analysis incorporating non-uniform temporal weighting of accumulated evidence (GLM4), in addition to the nuisance terms, we included the following regressors: |Δ*SV*|, lagged |Δ*AVnon-uniform|*, |Δ*SVGaze*| and lagged |Δ*AVnon-uniform, Gaze|*.

None of the regressors in the models were orthogonalized.

#### Region of interest specifications

ROIs were based upon previously published brain atlas parcellations and relevant literature. We used the Harvard-Oxford atlas for the intraparietal sulcus (IPS) and striatum (Desikan et al., 2006). The dorsolateral prefrontal cortex (dlPFC) and pre-supplementary motor area (pre-SMA) were defined based on (Hare et al., 2011). The ventromedial prefrontal cortex (vmPFC) was defined in (Bartra et al., 2013).

## Data Availability

Experiment and analysis code as well as Behavioral and eye-tracking data are available on the Open Science Framework: https://osf.io/eyxvb/files/

fMRI data is available on Open Neuro: doi:10.18112/openneuro.ds007305.v1.0.0

## Acknowledgements

Thanks to Kareem Soliman for research assistance with recruiting and to the Cattell Sabbatical Fund for financial support.

